# Protein Sequence Design by Entropy-based Iterative Refinement

**DOI:** 10.1101/2023.02.04.527099

**Authors:** Xinyi Zhou, Guangyong Chen, Junjie Ye, Ercheng Wang, Jun Zhang, Cong Mao, Zhanwei Li, Jianye Hao, Xingxu Huang, Jin Tang, Pheng Ann Heng

## Abstract

Inverse Protein Folding (IPF) is an important task of protein design, which aims to design sequences compatible with a given backbone structure. Despite the prosperous development of algorithms for this task, existing methods tend to leverage limited and noisy residue environment when generating sequences. In this paper, we develop an iterative sequence refinement pipeline, which can refine the sequence generated by existing sequence design models. It selects and retains reliable predictions based on the model’s confidence in predicted distributions, and decodes the residue type based on a partially visible environment. The proposed scheme can consistently improve the performance of a number of IPF models on several sequence design benchmarks, and increase sequence recovery of the SOTA model by up to 10%. We finally show that the proposed model can be applied to redesign Transposon-associated transposase B. 8 variants exhibit improved gene editing activity among the 20 variants we proposed. Our code and a demo of the refinement pipeline are provided in the online colab.

## 1 Introduction

Computational Protein Design, which is to design proteins with specific structures or functions [1], has been a powerful tool to prompt the exploration of sequence or topology space not yet visited by evolutionary process [2–4] and discover proteins with better properties [5]. It has enabled success in membrane protein design [6], enzyme design [7], etc. As one of the sub-tasks of Computational Protein Design, Inverse Protein Folding (IPF), the problem of finding amino acid sequences that can fold into a given three-dimensional (3D) structure [8], is of great importance as hosting a particular function often presupposes acquiring a specific backbone structure.

Traditional methods for IPF design energy functions to predict the compatibility between a sequence and the target backbone, which are usually linear combination of energy terms including force-field energies, solvation energies, etc [9, 10]. Residue-pair interaction modeling is usually derived from database by leveraging statistical preferences for particular residue pairs in the local environment to estimate inter-residue energies [5, 11, 12]. However, the residue interaction modeling in existing energy functions is restricted to residue pairs and the local environment representation is simplified to combinations of inter-residue distances, local secondary structures, backbone torsion angles, etc. Statistical estimation of multi-residue interaction conditional on more fine-grained local environment representation is limited by the increasing computational complexity [12, 13].

In recent years, deep learning has been widely and successfully applied to protein structure modeling and prediction [14, 15], due to its ability to automatically learn complex non-linear many-body interactions from data. There have been efforts to solve IPF by deep learning [4, 16, 17]. Early methods often model protein structures as sequences of independent residues [18, 19] or atom point clouds [4, 17] and adopt a non-autoregressive decoding scheme with all residues decoded in one shot, as demonstrated in Figure 1(1). Their independence assumption prevents them from learning complex residue inter-dependence and limits their performance. Some recent works adopt proximity graphs as protein structure representation with residues being nodes and residue interactions directly modeled as edges. A masked encoder-decoder architecture is generally employed [20–23], where the encoder first learns node representations based on local 3D structure only, and the decoder then recovers residue types from both structure representation and sequence information in an auto-regressive manner. However, the dependency on the previously generated sequence has proven to be prone to the error accumulation problem [24, 25]. Figure 1(2) demonstrates the auto-regressive decoding approach. Recently, ABACUS-R proposed in [26] also represents protein structures as graphs. Yet it assumes all neighbor residue types are known when decoding a central residue, as shown by Figure 1(3). It starts from a random initial sequence and recursively updates residue types based on their neighborhood until convergence. Similarly, the amino acid type decoding is conditioned on noisy prior predictions, which slows down the convergence to hundreds of iterations. However, recovering a target residue should be easier and more accurate if more ground truth neighbor types are available.

**Figure 1.**
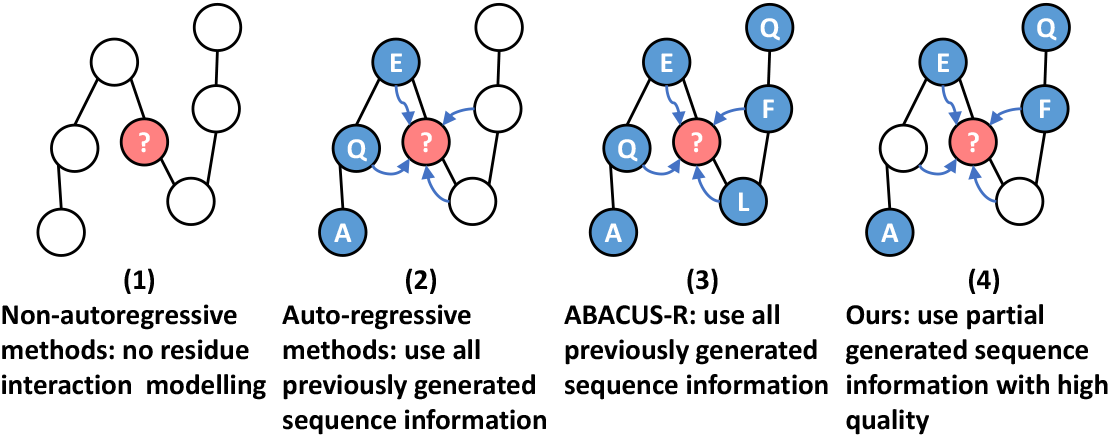
Different ways of utilizing inter-residue features. Our refinement model only leverages selected amino acid information with high quality and thus allows better residue recovery and faster convergence.

To address the above issues, we propose a residue sequence refinement scheme that can fully leverage the inter-residue features while alleviating error accumulation to refine a given generated sequence. The scheme is illustrated in Figure 2a. Specifically, a baseline model, which could be any existing model for IPF, first predicts an initial sequence in a form of probability distributions over 20 amino acids at every position. We select the residues that are likely to be decoded to ground truth types, retain their predicted types and discard others, resulting in a protein structure with partial residue type information. A proposed sequence refinement model then decodes a new sequence based on the partial environment. High quality predictions are selected likewise to supplement the partial environment, from which the refinement model makes new predictions again. The above procedure is iterated until convergence. Figure 1(4) illustrates the partial residue environment. To select the residues that are likely to be correctly recovered, we compute the entropy of the predicted distribution at each position and select the positions with low entropy, with the assumption that models are more confident with low-entropy predictions. In our experiment, the precision among the residues with the lowest 10% entropy is around 99%. Therefore, different from previous methods, our method can remove a large portion of noise in the input residue environment, which improves both the generated sequences and the converging speed. The final prediction will be the averaged prediction from every iteration weighted by their entropy. Our method has the advantage of being general and applicable to any existing IPF models as long as prediction entropy or other confidence measures could be computed.

**Figure 2.**
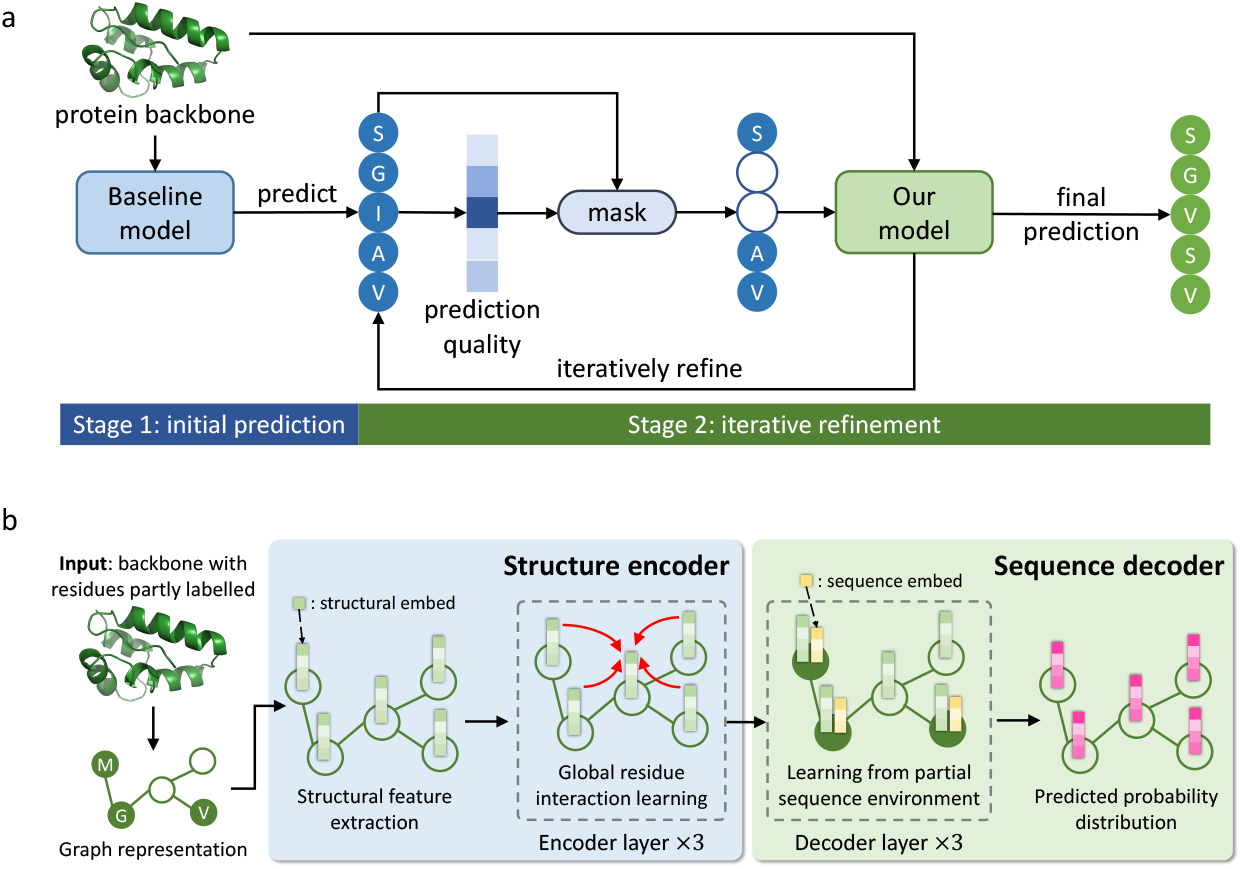
a. The sequence refinement scheme. The proposed refinement model repeatedly refines the sequence generated by a baseline model. In each iteration only high quality predictions are kept to reduce noise in the residue environment. b. The proposed refinement model. The structure encoder encodes structural embedding with global attention mechanism. The sequence decoder learns the sequence embedding from the input partial sequence, which is then aggregated with encoded structural embedding to predict a distribution over amino acid types.

Figure 2b shows the structure of the proposed sequence refinement model. The model input is the protein backbone structure represented as a graph, with part of (possibly no) residue nodes labeled with amino acid types. To better utilize the residue environment features, we design a global attention mechanism which can fully leverage graph edge features. Attention is originally designed for sequential data. In contrast to the localized nature of Graph Neural Networks (GNN), it provides a mechanism to model global dependency [27]. To adapt attention to graph domain, existing works either abandon the original global view [20, 28, 29] or require fully connected graphs to support global attention calculation [30, 31], which raises the memory complexity from linear to quadratic in node number. Besides, edge features are not fully leveraged since they only participate in attention calculation but cannot be updated or used to update node features [20, 28, 30, 32]. Yet we argue that edge features are essential in protein structure modeling, where multi-body interactions and most structural features, e.g. inter-residue distances, are represented on edges [22]. To address these issues, we construct a model that: (1) allows global residue attention, which enables modeling and learning of whole-structure features, (2) fully leverages edge features to better learn residue-pair interactions, (3) eliminates the need for fully-connected input graph, reducing the memory complexity to linear. Model design is detailed in Section 5.

We validate our sequence refinement method with different IPF models as the baseline model on several protein design benchmarks. Results show that our model can successfully refine the sequence generated by baseline models and boost the native sequence recovery rate. We also conduct ablation studies to verify the importance of the key designs in our model.

To further demonstrate our model’s ability to model the residue-environment interactions, we reinterpret our model’s output probability distribution at a certain position as a per-residue energy term describing the stability of this amino acid when the backbone and other amino acids are fixed. Then this model-predicted stability score could be used to assist in redesigning of proteins. We conduct experiments on Transposon-associated transposase B (TnpB), with the aim of finding variants with improved gene editing activity. Experiment results and details are presented in Section 2 and Section 5 respectively.

## 2 Results

### 2.1 Native Sequence Recovery

We train the proposed model on CATH 4.2 training set [20]. Training details are explained in Section 5. To evaluate our method, we take the following models as baseline model:

**GVP-GNN**[21] trained on the training split of CATH 4.2 containing 18,204 structures,

**ProteinMPNN**[22] trained on selected PDB structures clustered into 25,361 clusters, and the same model which we train on the CATH 4.2 training set for fair comparison and denote as **ProteinMPNN-C**,

**ESM-IF1**[23] trained on CATH 4.3 training set with 16,153 structures and 12 million additional structures predicted by Alphafold2 [15]. We conduct experiments on the following 5 benchmarks.

**CATH**. CATH 4.2 dataset [20] is a standard dataset for IPF training and evaluation. We evaluate on its test split of 1,120 structures.

**TS50** TS50 is a benchmark set of 50 protein chains proposed by [19]. It has been used by a number of previous works [17, 33, 34].

**EnzBench**EnzBench is a standard sequence recovery benchmark consisting of 51 proteins [35]. Designing algorithms are required to recover the native residues on the protein design shells. This benchmark is designed to test the algorithm’s ability to model protein binding and overall stability.

**BR EnzBench** BR EnzBench [36] aims to test the algorithm’s ability to remodel the chosen protein structure. It randomly selects 16 proteins from EnzBench benchmark and uses the Backrub server [37] to create an ensemble of 20 near-native conformations for each protein. To further increase the designing difficulty, all residues on the design shell are mutated to alanine, and conformations are then energy-minimized. When evaluated on EnzBench and BR EnzBench, types of residues not on design shells are fixed and available to models. Models can decode the design shell sequence conditioned on both structural information and residue environment.

**Latest PDB** We collect the latest published structures in PDB as another benchmark to validate the model’s ability to generalize to new structures. We select protein structures released after 01/01/2022 with a single chain of length less than 500 and resolution < 2.5 *Å*, which results in 1,975 protein structures.

We report two metrics on all benchmarks: sequence recovery and native sequence similarity recovery (nssr) [36]. A pair of residues is considered similar and contributes to the nssr score if their BLOSUM62 score [38] > 0. Compared with recovery which only considers residue identity, nssr takes residue similarity into account and provides a more specific comparison between two sequences. In Table 1, we report the median recovery rate and nssr scores for original and refined models, which we will denote with suffix’-R’ for simplicity. Among the original models, ESM-IF1 largely outperforms other models, indicating the effectiveness of data augmentation. On the two standard benchmarks CATH and TS50, the refinement scheme significantly improves both recovery and nssr for all models, demonstrating its ability to enhance residue relationship and similarity modeling. It also improves the baselines’ performance on EnzBench and BR EnzBench, which means it is also effective in cases where part of the sequence needs to be designed and the rest of it is fixed. Results on BR EnzBench further show its ability to remodel protein structures and recover native sequences even from corrupted near-native conformations. Similar results are also found on the Latest PDB benchmark. These experiments validate that our refinement approach is effective and applicable to various baseline models and designing scenarios.

**Table 1.**
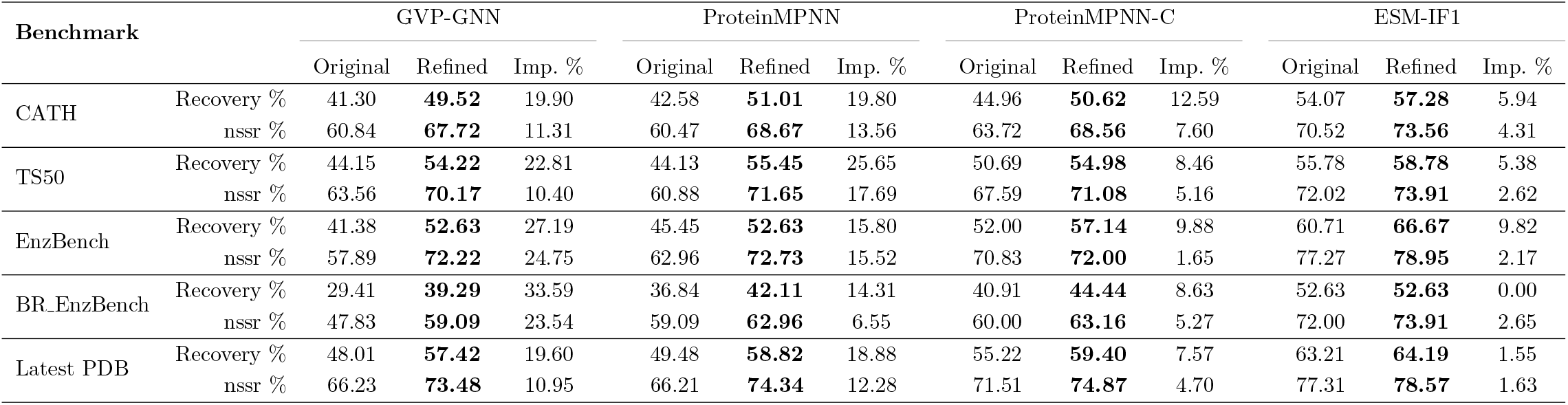
Median sequence recovery rate and nssr of different models on several benchmarks. “Original” and “Refined” denote the original models
and the corresponding models refined by our sequence refinement scheme respectively. Imp. shows the improvement rate.

We investigate the model performance on surface and core residues. In Figure 3a, we plot the recovery rate on CATH benchmark given different percentile in relative solvent accessible surface area (ASA). As expected, all models have higher recovery rate on core residues with low relative ASA since core residues are more constrained. The refinement operation leads to higher recovery over both core and surface residues, and brings slightly more benefits to recovering the surface residues, especially for ESM-IF1, where recovery for core residues is already high and the improvement of refinement is limited. This is further validated by Figure 3b, which shows the improvement rate on hydrophilic and hydrophobic residues. Except for GVP-GNN, other baseline models all have higher improvement rate for hydrophilic residues than hydrophobic ones. In Figure 4a, we break down the improvement rate to different secondary structures. In general, refinement helps the most in modeling *π*-helices and *α*-helices and is less helpful in recovering bends and turns.

**Figure 3.**
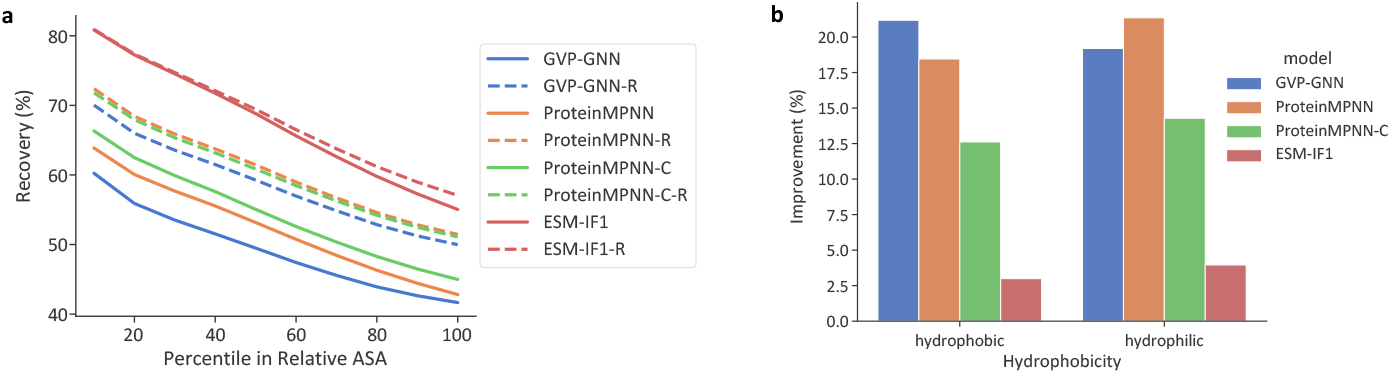
a. The native residue recovery of different models with respect to the percentile in relative ASA. b. The recovery improvement rate of our method for different baseline models over hydrophobic and hydrophilic residues.

**Figure 4.**
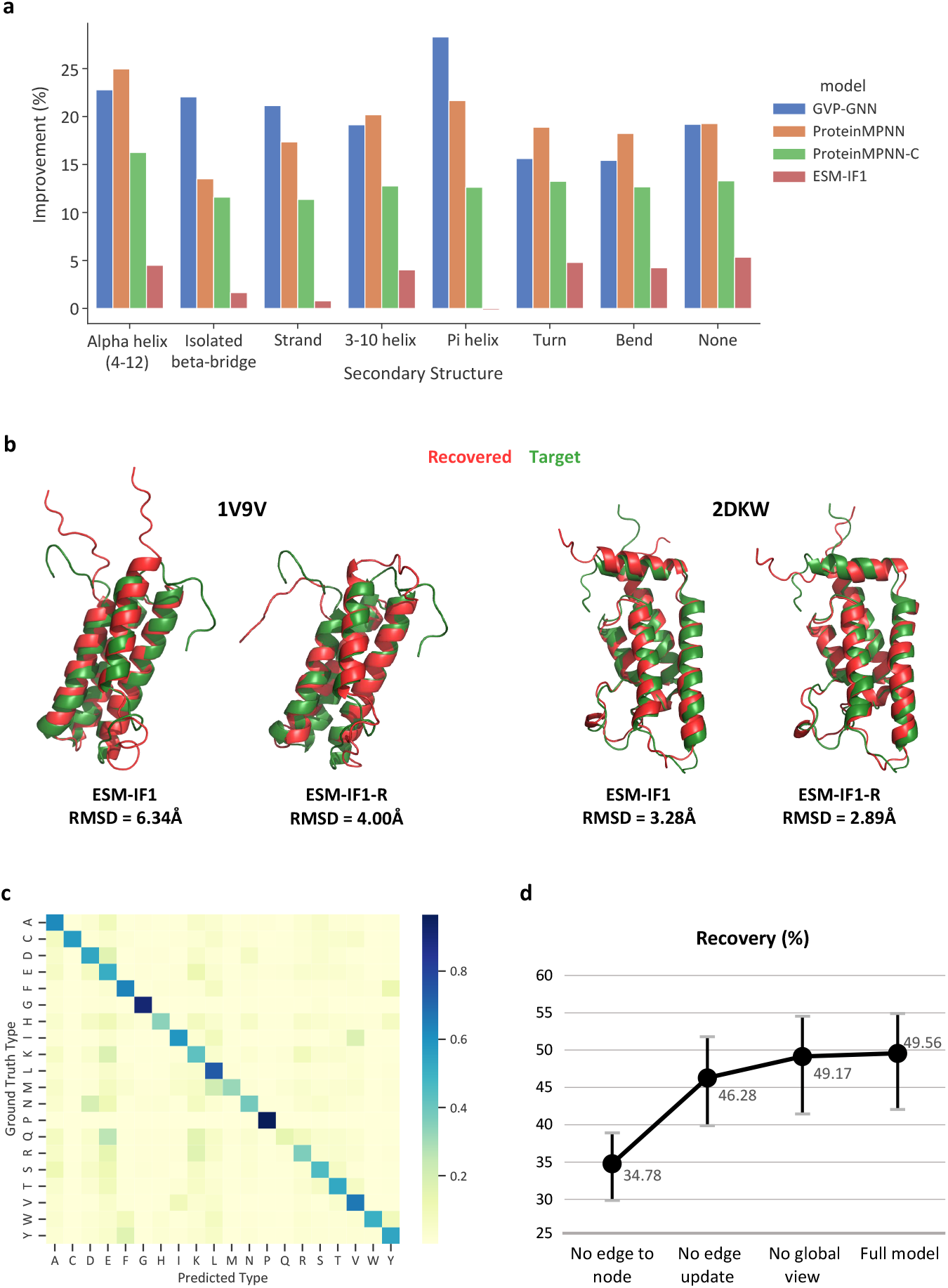
a. The recovery improvement rate of our method for baseline models over residues with different secondary structures. b. The Alphafold2 predicted structures of sequences recovered by ESM-IF1 and ESM-IF1-R compared to the target structures. c. The confusion matrix of ESM-IF1-R model. d. The recovery rate of ablated models.

To confirm if the recovered sequence can fold into the target structure, we compare the recovered structures predicted by Alphafold2 (in red) with the target native structures (in green) on 2 proteins from the CATH test set (Figure 4b). We focus on comparison with ESM-IF1 as it has the best performance. ESM-IF1-R has lower root-mean-square deviation (RMSD), which means the refined sequences are predicted to fold into structures with higher similarity to the target ones.

We calculate and show the confusion matrix of ESM-IF1-R, the best performing model, in Figure 4c. A darker cell means a larger portion of the residues of the native type (the vertical axis) is predicted to be the corresponding type on the horizontal axis. It can be observed that the residue types the model tends to confuse are also physicochemically similar types, such as ILE vs VAL and GLU vs LYS. We further estimate the correlation between the processed confusion matrix (following [17]) and the BLOSUM62 amino acid substation matrices [38] by Mantel test [39]. Results show that the p-value (< 0.0001) is lower than the significance level alpha = 0.05, indicating the two matrices are highly correlated.

### 2.2 Ablation Study

To validate the effectiveness of the key components in our method, we make one of the following modifications to our method and re-train the model.

- No edge to node. Do not use edge features to update node features in both encoder and decoder. Edge features only participate in attention calculation.
- No edge update. Do not update edge features.
- No global view. Remove global node attention in the encoder. Nodes only attend to their local neighbors.

Figure 4d shows the performance of ablated models on CATH test set. To better observe the performance change of model itself, we report the recovery rate of the sequences designed by ablated models alone without the baseline model. To do this, the types of all residues are masked and set to unknown in the input graph and the model is called only once. Using edge features to update node features is essential according to the figure, which confirms that it is crucial to fully leverage edge features on graphs where most information is carried by edges. Only allowing edges to compute attention is, therefore, not enough. Eliminating edge feature update prevents model from learning and propagating the residue interactions. Restricting nodes to local views makes it hard to establish long-term links between residues and model global protein structure. Therefore, these modifications also result in a considerable decrease in model performance.

## 3 Application: Mutation Site Recommendation for Transposon-associated transposase B

Transposon-associated transposase B (TnpB) is considered to be an evolutionary precursor to CRISPR-Cas system effector protein [40]. TnpB (408 amino acids) in the *D.radiodurans* ISDra2 element has been demonstrated to function as a hypercompact programmable RNA-guided DNA endonuclease [41], and its miniature size suitable for adeno-associated virus-based delivery. However, TnpB exhibits moderate gene editing activity in mammalian cells, limiting its therapeutic application.

Here we reinterpret our model’s output probability distribution at a position as a per-residue energy term describing the stability of this amino acid when the backbone and other amino acids are fixed. For a target position, the predicted probability of a certain amino acid type indicates the stability when this position mutates to the corresponding type. We hypothesized that our model could provide candidate mutation sites to enhance the activity of TnpB. With the intuition that a more positive surface charge might improve activity, we restrict the mutation target to arginine (R), which is the most positively charged amino acid, and restrict the candidate mutation sites to surface residues. Then we leverage the model prediction to rank the residue positions by their predicted stability.

Specifically, the native protein structure is first predicted by AlphaFold2 [15]. For each candidate position, we mask its residue type on the sequence. The sequence with one position masked and the predicted structure are fed into the model. The predicted probability for R on the target position is taken as the mutant stability score. A higher probability implies a higher compatibility between R and the given context and thus higher stability. Besides, based on empirical experience that mutation sites close to the TnpB binding site are more likely to bring improvements, we consider the distance between the *C_α_* of target position and the center of the predicted binding site (we leverage GraphSite [42] for binding site prediction). The stability score plus the negatives of the distance is taken as a quality score measuring how likely a mutation site can yield a stable and improved mutant. Figure 5a illustrates the process of calculating the quality score of one mutation site. All candidate sites are ranked in descending order and the first 20 are taken as our recommended mutation points.

**Figure 5.**
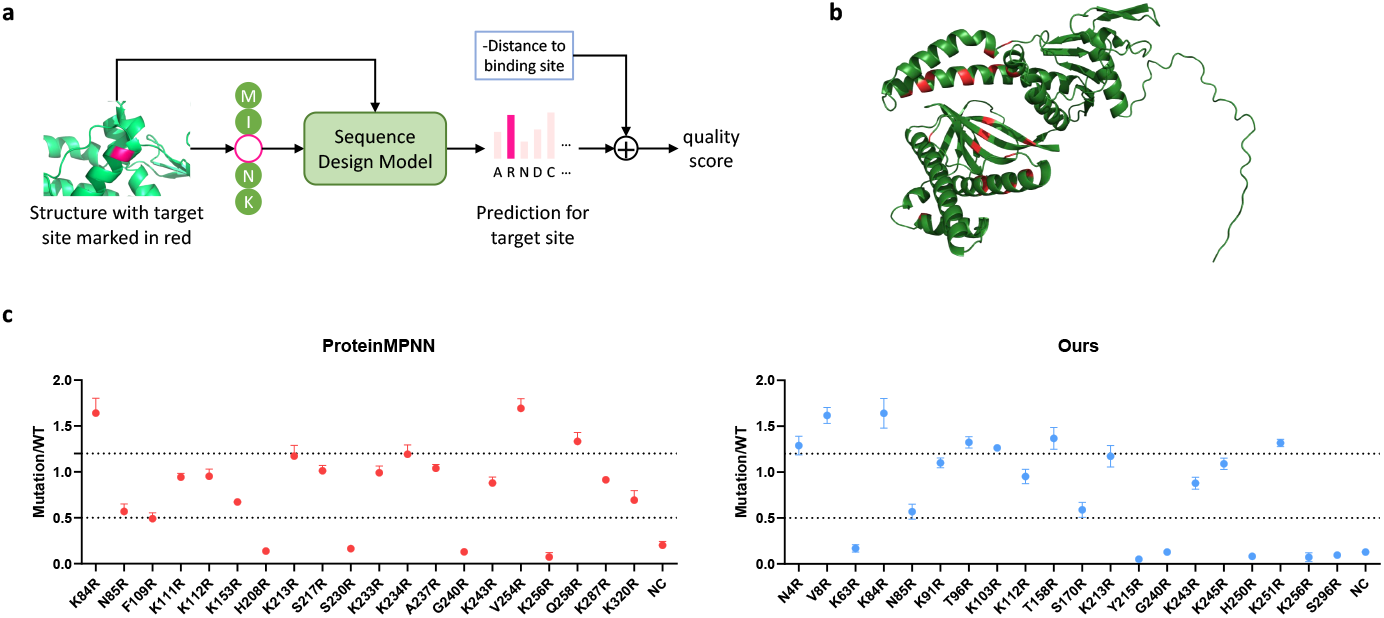
a. The process of computing the quality score of one target mutation site using our model. b. The mutation points recommended by our model marked in red. c. The improvement of variants recommended by ProteinMPNN and our model in indel activity relative to TnpB WT.

To test TnpB variants activity in human cells (HEK293T), plasmids encoding the TnpB variants fused with N-and C-terminal nuclear localization (NLS) sequences and reRNA construct targeting a EMX1 site in human genomic DNA (gDNA) were transiently transfected into HEK293T cells. After 96h, gDNA was extracted and analysed by sequencing for the presence of insertions and deletions (Indels) at the targeted cleavage sites. In Figure 5b, the 20 predicted sites are marked in red on TnpB predicted structure. Among them, 8 arginine substitutions showed above 1.2-fold improvement in indel activity relative to TnpB WT. ProteinMPNN is also employed to recommend mutation sites following the same procedure. For fair comparison, we use the ProteinMPNN trained on CATH training set (ProteinMPNN-C). With the stability score modeled by ProteinMPNN, 3 arginine substitutions with enhanced efficiency are found for TnpB. Results are given in Figure 5c.

This experiment verifies that our model can effectively model the residue interactions within a structural environment and predict the amino acid types that best fit a given 3D context. It can be leveraged to redesign existing proteins to improve stability or other qualities that highly depend on protein stability when used in conjunction with other property measures.

## 4 Conclusion

We develop a sequence refinement scheme that can refine the sequences generated by existing IPF models. We use entropy to select and keep reliable predictions and generate new sequences iteratively. A sequence refinement model is proposed to decode amino acids based on the 3D protein structure and partial sequence environment. Different from previous attention-based graph neural networks, the proposed model provides global node attention and fully leverages edge features while being memory-efficient. We argue that these features are not always guaranteed by existing works, yet they are essential for protein structure modeling, which is verified by our ablation studies. The effectiveness of our method is validated on several benchmarks with a number of recent IPF models as the baseline model.

We demonstrate one application of our model by redesigning TnpB. By leveraging the predicted probabilities to approximate protein stability, we are able to discover mutants with enhanced activity.

## 5 Methods

In this section, we present our method in detail. In Section 5.1-5.4, we provide the implementation of the proposed model and sequence refinement pipeline. In Section 5.5, we elaborate on the details of TnpB mutation site recommendation experiment.

### 5.1 Graph Representation of Protein Structures

A protein structure is represented as a proximity graph 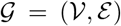, where 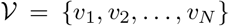, denotes the residue nodes and E = {*e_ij_*} denotes the directed edges form *v_j_* to *v_i_*, where residue *v_j_* is among the *k* = 30 nearest neighbors of *v_i_* in terms of *C_α_* distance. Each node *v_i_* has the following features:

- sin and cos value of dihedral angles;
- unit vectors from the previous and next residues on sequence to *v_i_* in terms of *C_α_* position.

Each edge *e_ij_* has the following features:

- Gaussian radial basis functions encoding of interatomic distances between *N, C_α_, C, O* and a virtual *C_β_*, and encoding of distance on sequence *i*−*j* [22];
- unit vector from *v_j_* to *v_i_* in terms of *C_α_* position.

We then employ 2 geometric vector perceptrons layers [21] to embed the extracted features to *d* dimension. Resulting node features are denoted as *H*° ∈ **R***^N ×d^* where 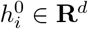 denotes the feature of *v_i_*. Resulting edge features are denoted as *E*° ∈ **R***^N ×k×d^* where 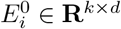 is the features of *k* neighbors of *v_i_* and 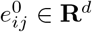 denotes the feature of edge from *v_j_* to *v_i_*.

### 5.2 Model Architecture

The proposed model adopts an encoder-decoder architecture. The encoder has a Transformer-based structure and produces encoded graph features. The decoder decodes residue type from a partially masked environment. Figure 2b demonstrates the model architecture.

#### 5.2.1 Encoder

Attention is first introduced in Transformer model [27]. Let *H* ∈ **R**^*N×d*^ denotes the *d*-dimension features of input sequence with length *N*. The self-attention module updates the input features according to the following equations:

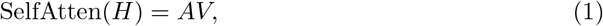

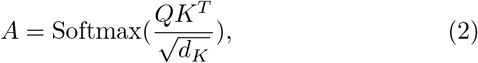

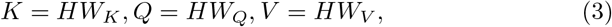

where 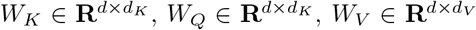 are parameters to map *H* to keys, queries and values.

There have been attempts to employ Transformer architecture for learning on graphs, with nodes denoted as sequence tokens. To utilize the global view provided by the original self-attention on graphs with edge features, previous works generally incorporate edge features into the attention matrix *A*:

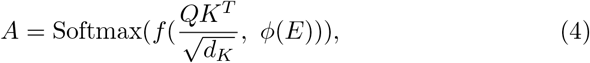

where *E* ∈ **R**^*N ×N ×d*^ is the the *d*-dimension edge features between each pair of nodes, *ϕ* estimates the correlations of node pairs from edge features, which could be linear transformation [30] or more sophisticated functions [32], and *f* is an aggregation function, which could be element-wise addition [30, 32] or multiplication [30]. These methods have two limitations. First, to construct the edge feature matrix *E*, they require fully connected graphs as input and the memory complexity will be 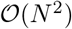. Second, the edge features are not fully leveraged. They are only involved in attention computation and can not be used to update node features or vice versa.

Our encoder is composed of a stack of *L* self-attention layers. In each layer, nodes can globally attend to all other nodes and edge features between node pairs serve as additive attention bias term. For non-existing edges, one solution is to convert arbitrary graphs to fully connected graphs before entering the encoder, then Equation 4 could be used. This could be done by setting *k* to a large enough number or using a fixed masking value/vector for non-existing edges as in previous works [30]. This operation increases memory complexity from 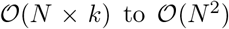. To avoid the conversion, we design a learnable pseudo edge feature in each self-attention layer. Let *H^l^* and *E^l^* denote the input features of the *l*th layer and *H*°, *E*° are inputs of the first layer. The attention is computed as follows:

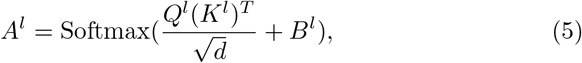

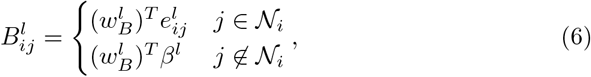

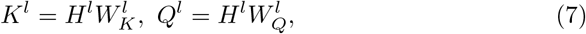

where *B^l^* is the attention bias, 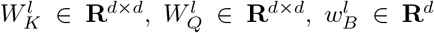 are parameters, *β^l^* ∈ **R**^*d*^ is the pseudo edge feature in layer *l*. Learning a pseudo edge feature for each layer is more adaptive and flexible than using one fixed masking value across all layers and provides a better approximation to using fully connected graphs.

The attention score *A^l^* is then used to aggregate node features as well as edge features. Node features are weighted and summed as in vanilla Transformer. Edge features are aggregated with normalized weights and concatenated with aggregated node features. Finally, a linear layer is employed to map the concatenated feature to dimension *d*:

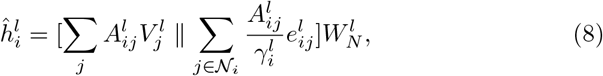

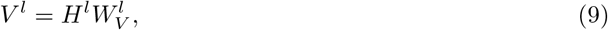

where 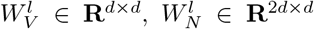 are parameters, ∥ means concatenation operation and 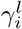 is a normalization term to normalize the sum of edge weights to 1.

Then a residue connection and layer normalization are adopted to output the final updated node features:

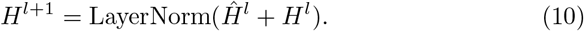

The edge features will then be updated as follows, with 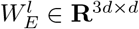 being parameters:

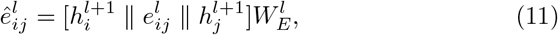

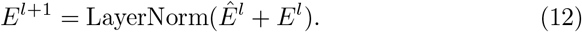

To leverage edge features under global attention mechanism, compared with 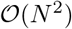 by previous works, our Transformer-based encoder only needs 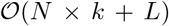 additional memory, and therefore allows designing longer sequences. Figure 6 compares the memory complexity with and without pseudo edge features. With pseudo edge features, the model runs in a linear memory complexity to input length. Removing it results in quadratic complexity.

**Figure 6.**
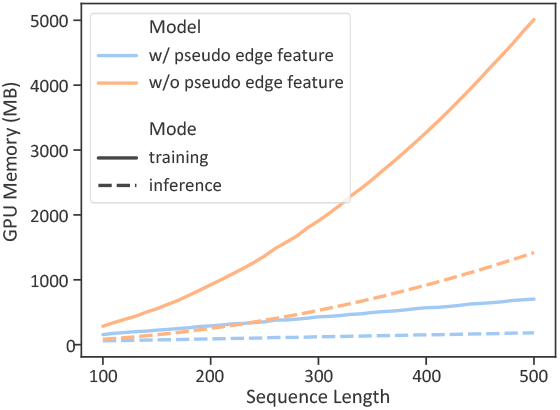
Memory consumption with and without pseudo edge feature.

#### 5.2.2 Decoder

The decoder is composed of a stack of *L′* decoder layers. We adopt the decoder framework from [22], with additional techniques of random partial masking and sequence noise. Let *H^l^*^(*dec*)^, *E^l^*^(*dec*)^ denote the input features for the *l*th layers and *H*^(*enc*)^, *E*^(*enc*)^ denote the output of encoder, then *H*^0(*dec*)^ = *H*^(*enc*)^.

**Random partial masking** For a graph with *N* residue nodes, we randomly sample a node subset 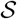 and set their ground truth residue type to be visible during training. This allows us to model not only the residue distribution conditioned on 3D structures but also the interaction between different types of residues in a more explicit way.

**Sequence noise** Among the type-visible nodes in 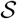, we further sample a smaller subset 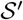 and set their type to a random type. We use *t_i_* to denote the type of 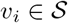 after applying the sequence noise.

In the *l*th decoder layer, the input edge features are constructed as follows:

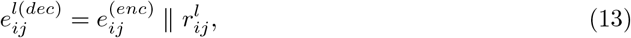

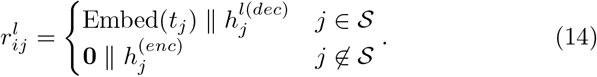

Then the node features will be updated as follows:

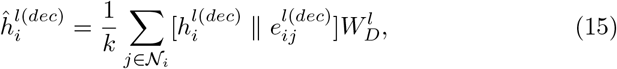

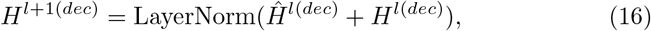

where 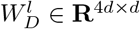 is the parameter. The output node features from the last layer will be mapped to the distribution over 20 residue types through a linear layer with parameter *W_P_* ∈ **R**^*d×*20^:

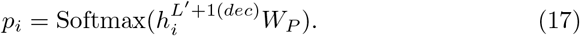

Negative log likelihood loss is used during training. In addition, we employ the DropEdge technique from [43] to prevent overfitting.

### 5.3 Sequence Refinement Pipeline

We design a sequence refinement pipeline, where a baseline model first gives an initial prediction, and our proposed model described in Section 5.2 will iteratively improve the sequence, as shown in Figure 2a.

Suppose 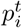 is the probability distribution of *v_i_* predicted in iteration *t*, with 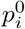 given by the baseline model. We compute the entropy 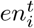 of distribution 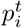:

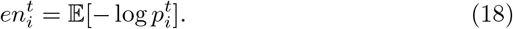

*th × N* nodes (0 <*th*< 1) with the least entropy are selected and added to the visible node set in iteration *t* + 1 as the nodes in 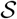 during training. We stop iteration when few new residues are selected. Finally, the predicted distributions from all iterations will be fused as follows,

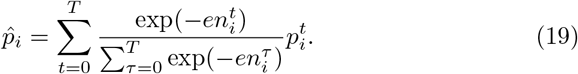

*T* is the total number of iterations. Final predicted residue type will be the argmax of 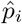.

### 5.4 Experiment Settings

We use *L* = 3 encoder layers with 8 attention heads and *L′* = 3 decoder layers. We train our model on CATH 4.2 training set. During training, we set the size of visible residue set 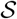 as *N*/10 and sample 3% from them to apply type noise. We use dropout rate 0.2 and drop edge rate 0.3. We also apply multi-task training and let the model predict the secondary structure type, relative accessible surface area and backbone torsional angles (*ϕ, Ψ*) together with amino acid type for every residue. Adam optimizer [44] is employed with learning rate 0.001 and cosine annealing scheduler [45]. During inference, the threshold *th* is selected based on sequence recovery on CATH 4.2 validation set for each baseline model.

### 5.5 Experiment Details of TnpB Variant Recommendation

#### Plasmid vector construction

The TnpB gene was optimized for expression in human cells through codon optimization and the optimized sequence was synthesized for vector construction (Sangon Biotech). The final optimized sequence was inserted into a pST1374 vector, which contained a CMV promoter and nuclear localization signal sequences at both the 5 *^′^* and 3*^′^* termini. reRNA sequences were synthesized and cloned into a pGL3-U6 vector. Spacer sequences (EMX1: 5^*′*^-ctgtttctcaggatgtttgg −3^*′*^) were cloned into by digesting the vectors with BsaI restriction enzyme (New England BioLabs) for 2 h at 37*^°^C*. The resulting vector constructs were verified through Sanger sequencing to ensure accuracy.

#### TnpB engineering

The construction of TnpB mutants was carried out through the use of site-directed mutagenesis. PCR amplifications were performed using Phanta Max Super-Fidelity DNA Polymerase (Vazyme). The PCR products were then ligated using 2X MultiF Seamless Assembly Mix (ABclonal). Ligated products were transformed into DH5*α* E. coli cells. The success of the mutagenesis was confirmed through Sanger sequencing. The modified plasmid vectors were purified using a TIANpure Midi Plasmid Kit (TIANGEN).

#### Cell culture and transfection

HEK293T cells were maintained in Dulbecco’s modified Eagle medium (Gibco) supplemented with 10% fetal bovine serum (Gemini) and 1% penicillin–streptomycin (Gibco) in an incubator (37*^°^C*, 5% CO_2_). HEK293T cells were transfected at 80% confluency with a density of approximately 1 × 10^5^ cells per 24-well using ExFect Transfection Reagent (Vazyme). For indel analysis, 500 ng of TnpB plasmid plus 500 ng of reRNA plasmid was transfected into 24-well cells.

#### DNA extraction and Deep sequencing

The genomic DNA of HEK293T cells was extracted using QuickExtract DNA Extraction Solution (Lucigen). Samples were incubated at 65 *^°^C* for 60 minutes and 98 *^°^C* for 2 minutes. The lysate was used as a PCR template. The first round PCR (PCR1) was conducted with barcoded primers to amplify the genomic region of interest using Phanta Max Super-Fidelity DNA Polymerase (Vazyme). The products of PCR1 were pooled in equal moles and purified for the second round of PCR (PCR2). The PCR2 products were amplified using index primers (Vazyme) and purified by gel extraction for sequencing on the Illumina NovaSeq platform. The specific barcoded primers used in PCR1 are listed in Table 2.

**Table 2.**
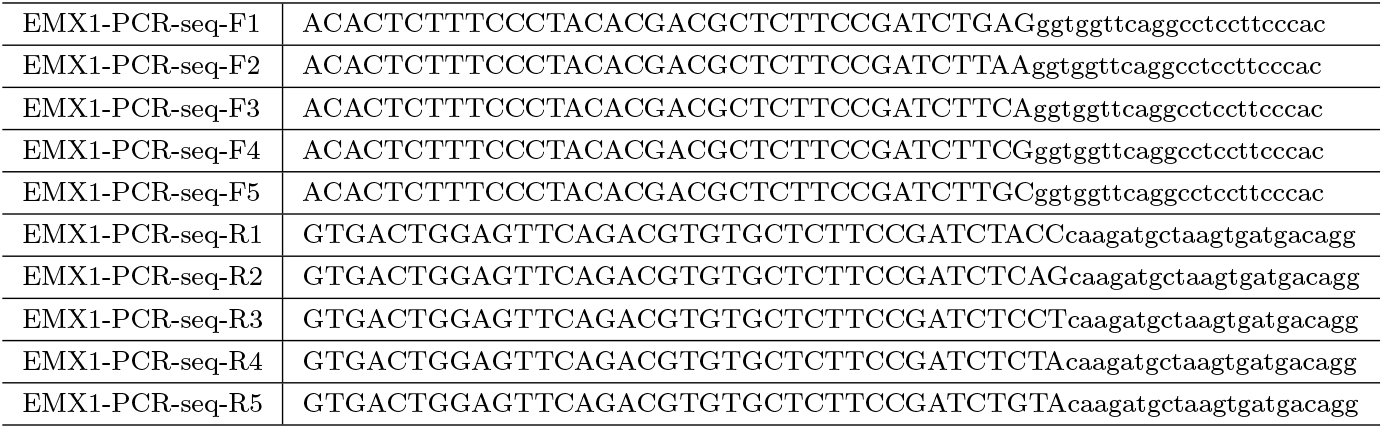
The barcoded primers used in PCR1.

## 6 Acknowledgement

This work was supported by National Key R&D Program of China (2022YFE0200700, 2022YFC2702705), Hong Kong Innovation and Technology Fund (ITS/170/20 and ITS/241/21), National Natural Science Foundation of China (62006219, 22007082), Natural Science Foundation of Guangdong Province (2022A1515011579), and Natural Science Foundation of Zhejiang Province, China (LQ21B030013).

## Notes

### Competing Interest Statement

The authors have declared no competing interest.

## References

[1] Gao W, Mahajan SP, Sulam J, Gray JJ. Deep learning in protein structural modeling and design. Patterns. 2020;p. 100142.

[2] Huang PS, Feldmeier K, Parmeggiani F, Fernandez Velasco DA, Höocker B, Baker D. De novo design of a four-fold symmetric TIM-barrel protein with atomic-level accuracy. Nature chemical biology. 2016;12(1):29–34.

[3] Lin YR, Koga N, Tatsumi-Koga R, Liu G, Clouser AF, Montelione GT, et al. Control over overall shape and size in de novo designed proteins. Proceedings of the National Academy of Sciences. 2015;112(40):E5478–E5485.

[4] Anand-Achim N, Eguchi RR, Mathews II, Perez CP, Derry A, Altman RB, et al. Protein sequence design with a learned potential. bioRxiv. 2021;p. 2020–01.

[5] Alford RF, Leaver-Fay A, Jeliazkov JR, O’Meara MJ, DiMaio FP, Park H, et al. The Rosetta all-atom energy function for macromolecular modeling and design. Journal of chemical theory and computation. 2017;13(6):3031–3048.

[6] Slovic AM, Summa CM, Lear JD, DeGrado WF. Computational design of a water-soluble analog of phospholamban. Protein Science. 2003;12(2):337–348.

[7] Jiang L, Althoff EA, Clemente FR, Doyle L, Röothlisberger D, Zanghellini A, et al. De novo computational design of retro-aldol enzymes. science. 2008;319(5868):1387–1391.

[8] Pabo C. Molecular technology: designing proteins and peptides. Nature. 1983;301(5897):200–200.

[9] Pokala N, Handel TM. Energy functions for protein design I: efficient and accurate continuum electrostatics and solvation. Protein science. 2004;13(4):925–936.

[10] Fan C, Yaoxi C, Yangyang M, Lu ZHANG HL. Computational protein design: perspectives in methods and applications. Synthetic Biology Journal. 2021;2(1):15.

[11] Wilmanns M, Eisenberg D. Three-dimensional profiles from residue-pair preferences: identification of sequences with beta/alpha-barrel fold. Proceedings of the National Academy of Sciences. 1993;90(4):1379–1383.

[12] Zhou X, Xiong P, Wang M, Ma R, Zhang J, Chen Q, et al. Proteins of well-defined structures can be designed without backbone readjustment by a statistical model. Journal of structural biology. 2016;196(3):350–357.

[13] Rohl CA, Strauss CE, Misura KM, Baker D. Protein structure prediction using Rosetta. In: Methods in enzymology. vol. 383. Elsevier; 2004. p. 66–93.

[14] Du Y, Meier J, Ma J, Fergus R, Rives A. Energy-based models for atomic-resolution protein conformations. arXiv preprint arXiv:200413167. 2020;.

[15] Jumper J, Evans R, Pritzel A, Green T, Figurnov M, Ronneberger O, et al. Highly accurate protein structure prediction with AlphaFold. Nature. 2021;596(7873):583–589.

[16] Norn C, Wicky BI, Juergens D, Liu S, Kim D, Koepnick B, et al. Protein sequence design by explicit energy landscape optimization. bioRxiv. 2020;.

[17] Zhang Y, Chen Y, Wang C, Lo CC, Liu X, Wu W, et al. ProDCoNN: Protein design using a convolutional neural network. Proteins: Structure, Function, and Bioinformatics. 2020;88(7):819–829.

[18] Li Z, Yang Y, Faraggi E, Zhan J, Zhou Y. Direct prediction of profiles of sequences compatible with a protein structure by neural networks with fragment-based local and energy-based nonlocal profiles. Proteins: Structure, Function, and Bioinformatics. 2014;82(10):2565–2573.

[19] O’Connell J, Li Z, Hanson J, Heffernan R, Lyons J, Paliwal K, et al. SPIN2: Predicting sequence profiles from protein structures using deep neural networks. Proteins: Structure, Function, and Bioinformatics. 2018;86(6):629–633.

[20] Ingraham J, Garg V, Barzilay R, Jaakkola T. Generative models for graph-based protein design. Advances in neural information processing systems. 2019;32.

[21] Jing B, Eismann S, Suriana P, Townshend RJ, Dror R. Learning from protein structure with geometric vector perceptrons. arXiv preprint arXiv:200901411. 2020;.

[22] Dauparas J, Anishchenko I, Bennett N, Bai H, Ragotte RJ, Milles LF, et al. Robust deep learning based protein sequence design using ProteinMPNN. bioRxiv. 2022;.

[23] Hsu C, Verkuil R, Liu J, Lin Z, Hie B, Sercu T, et al. Learning inverse folding from millions of predicted structures. bioRxiv.2022;.

[24] Li B, Tian J, Zhang Z, Feng H, Li X. Multitask non-autoregressive model for human motion prediction. IEEE Transactions on Image Processing. 2020;30:2562–2574.

[25] Huang R, Hu H, Wu W, Sawada K, Zhang M. Dance Revolution: Long Sequence Dance Generation with Music via Curriculum Learning. CoRR. 2020;abs/2006.06119.

[26] Liu Y, Zhang L, Wang W, Zhu M, Wang C, Li F, et al. Rotamer-free protein sequence design based on deep learning and self-consistency. Nature Computational Science. 2022 Jul;2(7):451–462.

[27] Vaswani A, Shazeer N, Parmar N, Uszkoreit J, Jones L, Gomez AN, et al. Attention is all you need. In: Advances in neural information processing systems; 2017. p. 5998–6008.

[28] Dwivedi VP, Bresson X. A Generalization of Transformer Networks to Graphs. CoRR. 2020;abs/2012.09699.

[29] Hu Z, Dong Y, Wang K, Sun Y. Heterogeneous Graph Transformer. In: Huang Y, King I, Liu T, van Steen M, editors. WWW’20: The Web Conference 2020, Taipei, Taiwan, April 20-24, 2020. ACM / IW3C2; 2020. p. 2704–2710.

[30] Hussain MS, Zaki MJ, Subramanian D. Edge-augmented Graph Transformers: Global Self-attention is Enough for Graphs. CoRR. 2021;abs/2108.03348.

[31] Bergen L, O’Donnell TJ, Bahdanau D. Systematic Generalization with Edge Transformers. In: Ranzato M, Beygelzimer A, Dauphin YN, Liang P, Vaughan JW, editors. Advances in Neural Information Processing Systems 34: Annual Conference on Neural Information Processing Systems 2021, NeurIPS 2021, December 6-14, 2021, virtual; 2021. p. 1390–1402.

[32] Ying C, Cai T, Luo S, Zheng S, Ke G, He D, et al. Do Transformers Really Perform Bad for Graph Representation? CoRR. 2021;abs/2106.05234.

[33] Wang J, Cao H, Zhang JZ, Qi Y. Computational protein design with deep learning neural networks. Scientific reports. 2018;8(1):1–9.

[34] Qi Y, Zhang JZ. DenseCPD: improving the accuracy of neural-network-based computational protein sequence design with DenseNet. Journal of chemical information and modeling. 2020;60(3):1245–1252.

[35] Nivóon LG, Bjelic S, King C, Baker D. Automating human intuition for protein design. Proteins: Structure, Function, and Bioinformatics. 2014;82(5):858–866.

[36] Löoffler P, Schmitz S, Hupfeld E, Sterner R, Merkl R. Rosetta: MSF: a modular framework for multi-state computational protein design. PLoS computational biology. 2017;13(6):e1005600.

[37] Lauck F, Smith CA, Friedland GF, Humphris EL, Kortemme T. Roset-taBackrub—a web server for flexible backbone protein structure modeling and design. Nucleic acids research. 2010;38(suppl 2):W569–W575.

[38] Henikoff S, Henikoff JG. Amino acid substitution matrices from protein blocks. Proceedings of the National Academy of Sciences. 1992;89(22):10915–10919.

[39] Mantel N. The detection of disease clustering and a generalized regression approach. Cancer research. 1967;27(2 Part 1):209–220.

[40] Makarova KS, Wolf YI, Iranzo J, Shmakov SA, Alkhnbashi OS, Brouns SJ, et al. Evolutionary classification of CRISPR–Cas systems: a burst of class 2 and derived variants. Nature Reviews Microbiology. 2020;18(2):67–83.

[41] Karvelis T, Druteika G, Bigelyte G, Budre K, Zedaveinyte R, Silanskas A, et al. Transposon-associated TnpB is a programmable RNA-guided DNA endonuclease. Nature. 2021;599(7886):692–696.

[42] Yuan Q, Chen S, Rao J, Zheng S, Zhao H, Yang Y. AlphaFold2-aware protein–DNA binding site prediction using graph transformer. Briefings in Bioinformatics. 2022;23(2):bbab564.

[43] Rong Y, Huang W, Xu T, Huang J. DropEdge: Towards Deep Graph Convolutional Networks on Node Classification. In: 8th International Conference on Learning Representations, ICLR 2020, Addis Ababa, Ethiopia, April 26-30, 2020. OpenReview.net; 2020.

[44] Kingma DP, Ba J. Adam: A Method for Stochastic Optimization. In: Bengio Y, LeCun Y, editors. 3rd International Conference on Learning Representations, ICLR 2015, San Diego, CA, USA, May 7-9, 2015, Conference Track Proceedings; 2015..

[45] Loshchilov I, Hutter F. SGDR: Stochastic Gradient Descent with Restarts. CoRR. 2016;abs/1608.03983.

